# Facilitation between ecosystem engineers, salt marsh grass and mussels, produces pattern formation on salt marsh shorelines

**DOI:** 10.1101/2021.04.14.439864

**Authors:** Romuald N. Lipcius, David G. Matthews, Leah Shaw, Junping Shi, Sofya Zaytseva

## Abstract

Interspecific facilitation between ecosystem engineers, such as salt marsh grass and mussel aggregations, is a key process that structures communities and enhances biodiversity. Scale-dependent pattern formation via self-organization is ubiquitous in terrestrial, aquatic and marine ecosystems. Despite their prevalence and ecological importance, these two phenomena have rarely been linked. We provide empirical evidence that the facilitative interaction in salt marshes between smooth cordgrass *Spartina alterniflora* and the ribbed mussel *Geukensia demissa* produces distinct spatial patterns along marsh shorelines. These findings advance our understanding of linkages between facilitation and pattern formation in nature, and are particularly relevant to conservation and restoration of salt marshes threatened by climate change and sea-level rise.

## Introduction

Self-organization and scale-dependent pattern formation (SDPF) are ubiquitous in terrestrial, aquatic and marine ecosystems, such as arid grasslands, savanna vegetation, wetlands, intertidal mussel beds, and coral reefs (*1–3*). In marine pelagic habitats, self-organization is a prevalent feature of species from the tropics to the poles, such as phytoplankton aggregations (*4*) and schooling fish (*5*). In marine benthic communities, the species involved in pattern formation include primary producers (e.g., benthic diatoms, salt marsh grasses, seagrasses) and invertebrate ecosystem engineers (e.g., mussels, corals and polychaete worms) (*6–14*). Geometric features of the seascape of ecosystem engineers can significantly influence community structure and diversity (*15*), such as in seagrass meadows (*16–19*), oyster reefs (*16, 17, 20, 21*) and mussel beds (*22*). Moreover, SDPF and positive interactions between species have recently been linked theoretically and experimentally (*23*) as a “landscape of facilitation” to determine their influence upon diversity and ecosystem function. Here we document a unique facilitative interaction between two ecosystem engineers, specifically salt marsh grass *Spartina alterniflora* and the ribbed mussel *Geukensia demissa*, that produces distinct spatial patterns along coastal shorelines.

## Materials & Methods

A mensurative field experiment was conducted along a salt marsh shoreline on the south shore of the York River, a western shore tributary of Chesapeake Bay. The shoreline consisted of marsh patches of different geometric patterns, including fingers, lobes, linear patches, and islands (Fig. 1). These were defined by two criteria: (i) length:width ratio (LWR), and (ii) whether or not the patch was connected to the shoreline. Length was measured outward from the shoreline and width parallel to the shoreline. ‘Islands’ were disconnected from the shoreline, while the remaining three patterns were contiguous with the shoreline. ‘Fingers’ had LWR *>*1, ‘lobes’ had LWR from 0.2 to 0.9 with a mean = 0.5, and linear patches had LWR *<*0.2. At the shoreline we randomly selected equal numbers of each pattern and measured mussel and cordgrass shoot density with a 13 cm x 13 cm quadrat placed on the leading edge of the patch.

**Figure 1:**
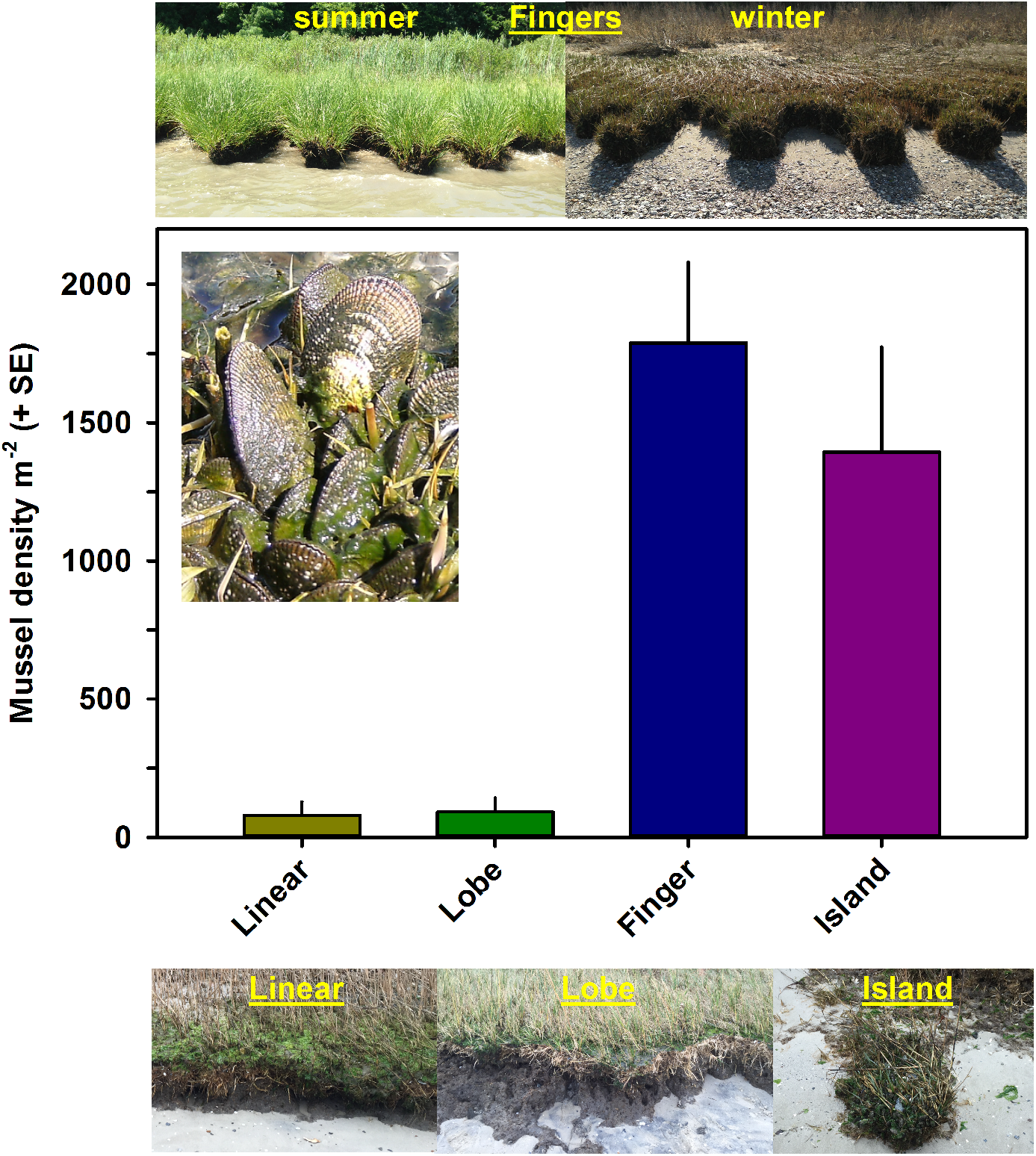
Mean mussel density at the four marsh types. All photos by R.N. Lipcius.

To analyze mussel density, we formed statistical models (*g*_*i*_) representing multiple alternative hypotheses (*24*), which were evaluated following an Information Theoretic approach (*25, 26*). The suite of models included the sparse and global models (Table 1). The response variable was density m^*-*2^, which was log_10_-transformed to meet statistical assumptions of linearity, normality and homogeneity of variance. The two independent variables were pattern type as a factor with four levels (island, finger, lobe and linear) and log_10_-transformed cordgrass shoot density as a continuous variable. The 5 alternative models included the sparse (null) model (*g*_1_), only marsh pattern type influencing density (*g*_2_), only shoot density (*g*_3_), both pattern type and shoot density in an additive model (*g*_4_), and pattern type and shoot density in a multiplicative model (*g*_5_).

**Table 1:** AIC calculations for the general linear models (*g*_*i*_) corresponding to the different hypotheses for log_10_-transformed mussel density on marsh front edge. Model *g*_4_ was significantly better than the null model *g*_1_ (Likelihood ratio *X*^2^, *Delta Deviance* = 48.80; *df* = 32, 28; *p <* 0.001), and the global model *g*_5_ did not reduce deviance significantly better than *g*_4_ (Likelihood ratio *X*^2^, Δ *Deviance* = -3.46; *df* = 25, 28; *p* = 0.07). df = degrees of freedom; k = number of parameters, including sample variance (*s*^2^) as a parameter; *AIC*_*c*_ = corrected AIC value; Δ_*i*_ = difference between model *g*_*i*_ and the best model (*g*_4_); *w*_*i*_ = model probability of fitting the observed data. Abbreviations: I = intercept; MPT = marsh pattern type; CSD = log_10_-transformed cordgrass shoot density.

| Model | Variables | $k$ | $AIC_c$ | $\Delta_i$ | $w_i$ |
| --- | --- | --- | --- | --- | --- |
| $g_1$ | I | 2 | 120.3 | 35.5 | <0.01 |
| $g_2$ | I + MPT | 5 | 90.0 | 5.2 | <0.01 |
| $g_3$ | I + CSD | 3 | 115.0 | 30.2 | <0.01 |
| $g_4$ | I + MPT + CSD | 6 | 84.8 | 0 | 0.73 |
| $g_5$ | I + (MPT x CSD) | 9 | 87.3 | 2.5 | 0.21 |

Each of the models was run in R using the glm procedure (*27*). The resulting Akaike Information Criterion (AIC) values from each model were used to calculate AICc, a second-order bias correction estimator (*26*). Model probabilities (*w*_*i*_), based on Δ_*i*_ values were used to rank the different models (*g*_*i*_) against the model with the lowest AICc, and estimated the probability that a particular model *g*_*i*_ was the best model. Following convention any model with *w*_*i*_ *<*0.10 was eliminated (*26*). Likelihood ratio *X*^2^ tests (*28*) were used to compare models.

## Results

Mean mussel densities ranged from nearly 1,800 m^*-*2^ on fingers and about 1,400 m^*-*2^ on islands to less than 100 m^*-*2^ on lobes and linear patches (Figure 1). Mussel density was best explained by the additive model (*g*_4_) of marsh pattern type and cordgrass shoot density, which had a weighted probability of 0.73, and which had a significantly better fit than all other models (Table 1). Model *g*_4_ explained 75.4% of the deviance and a multiple adjusted *r*^2^ = 71.9% (*F* = 21.47; *df* = 4, 28; *p <* 0.001). Mussel density was highest on fingers (Figure 1) but did not differ significantly from density on islands (Table 3). Mussel densities on fingers and islands were significantly higher than densities on lobes and linear patches, which did not differ significantly from each other (Figure 1, Table 3). Mussel density was also positively and significantly correlated with cordgrass shoot density on all marsh pattern types (Figure 2, Table 2).

**Table 2:** Parameter estimates from general linear model *g*_4_ for mussel density. Deviance explained = 75.4%. Multiple adjusted *r*^2^ = 71.9% (*F* = 21.47; *df* = 4, 28; *p <* 0.001). Variance in mussel density was not heterogeneous according to marsh pattern type (Levene’s test, *F* = 1.52; *df* = 3, 29; *p* = 0.23).

| Parameter | Estimate | SE | $p$ |
| --- | --- | --- | --- |
| Intercept (Finger) | -2.45 | 2.04 | 0.24 |
| Linear | -2.14 | 0.41 | < 0.001 |
| Lobe | -2.56 | 0.36 | < 0.001 |
| Island | -0.78 | 0.41 | 0.07 |
| CSD (slope) | 2.06 | 0.73 | 0.009 |

**Table 3:** Multiple comparison tests (Tukey) of marsh pattern types for model *g*_4_.

| Contrast | Estimate | SE | $p$ |
| --- | --- | --- | --- |
| Linear - Finger | -2.14 | 0.41 | <0.001 |
| Island - Finger | -0.78 | 0.41 | 0.23 |
| Lobe - Finger | -2.56 | 0.36 | <0.001 |
| Island - Linear | 1.36 | 0.51 | 0.04 |
| Lobe - Linear | -0.43 | 0.43 | 0.75 |
| Lobe - Island | -1.78 | 0.39 | <0.001 |

**Figure 2:**
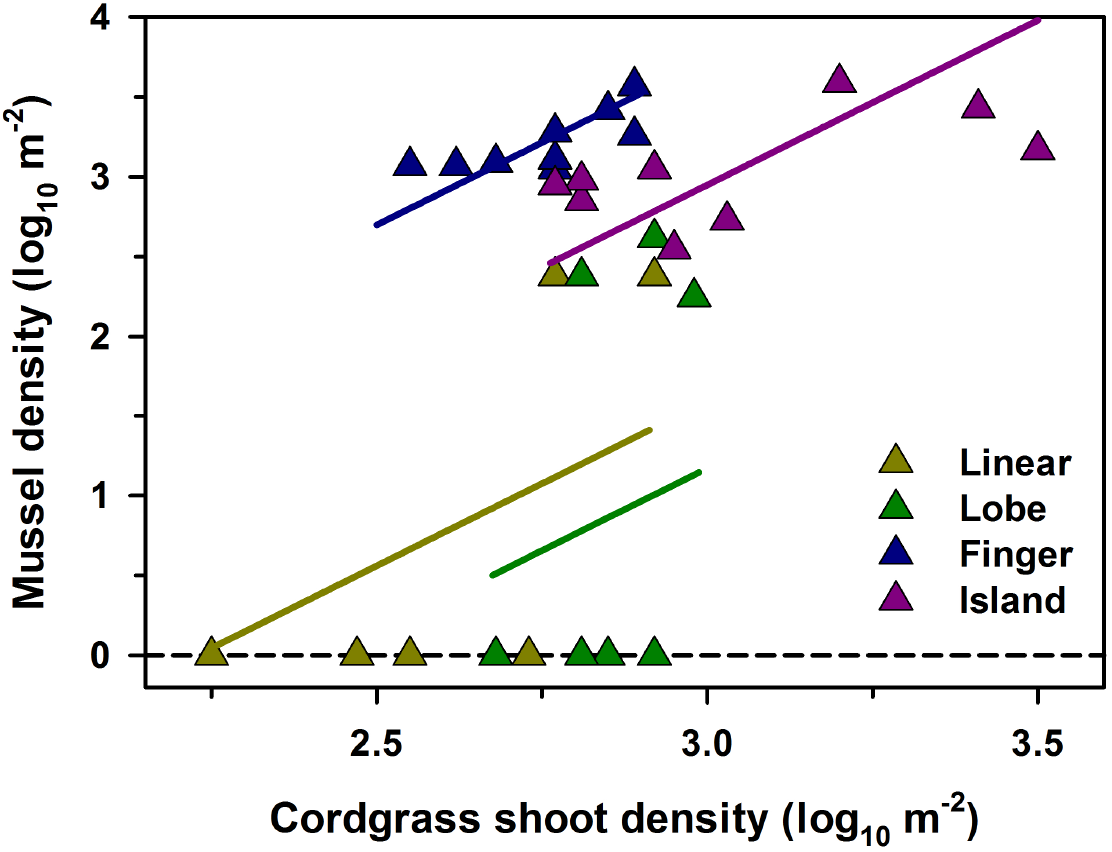
Mussel density as a function of cordgrass density at the four marsh types. The lines are derived from the parameter estimates of model *g*_4_, and only span the range of *x* values for each marsh type.

## Discussion

The novelty of the findings from our mensurative experiment is that the geometry and pattern formation of salt marsh shorelines is produced by interspecific facilitation of two dominant ecosystem engineers, ribbed marsh mussels and salt marsh cordgrass. Interspecific facilitation between marsh mussels and cordgrass has been extensively documented (*29,30*), but not pattern formation. Specifically, high densities of both species on a single patch produced long, narrow finger-like marsh projections bounded on both sides by eroded shorelines with low mussel and cordgrass densities. Similarly, marsh islands disconnected from the shoreline had high mussel and cordgrass densities, but these were the remnants of marsh fingers that had been eroded near the base where mussel and cordgrass densities were low. Hence, the marsh fingers reflect a transient marsh pattern that eventually erodes to an island pattern with the base of the finger becoming a lobe or linear patch. In contrast, marsh lobes and linear shorelines are characterized by low mussel and cordgrass densities. A further difference between the marsh patterns is that fingers occur at wavelenghts of a few meters whereas lobes and linear patches occur at much longer wavelengths ranging from several meters to 10s of meters (unpublished data).

We posit that the mechanisms driving scale-dependent pattern formation (SDPF) in salt marsh ecosystems display consistency with those producing SPDF in other marine ecosystems. In marine ecosystems, both short-range facilitation and long-range inhibition are driven by some combination of population density, resource availability, habitat quality, consumer pressure, and physical stress. Short-range facilitation can result from resource concentration (e.g., nutrients), substrate stability (e.g., sediment binding in mud flats), or enhanced survival at high density (e.g., reduced predation and physical stress in mussel beds) within habitat patches. Long-range inhibition is a consequence of resource depletion, substrate instability, or decreased survival due to physical or biotic factors (e.g., heavy siltation) away from habitat patches. In some cases, consumers can either accentuate (e.g., waterfowl feeding on seagrass) or obliterate (e.g., amphipods feeding on diatoms) patterns. Similarly, physical forcing such as tidal currents over intertidal flats can either alter or disrupt SDPF (*31*).

## Notes

### Competing Interest Statement

The authors have declared no competing interest.

## References

1. S. Camazine, Natural History 112, 34 (2003).

2. R. Solé, J. Bascompte, Self-Organization in Complex Ecosystems, vol. 42 (Princeton University Press, 2006).

3. M. Rietkerk, J. Van de Koppel, Trends in Ecology & Evolution 23, 169 (2008).

4. J. H. Steele, Spatial Pattern in Plankton Communities (Springer, 1978), pp. 1–20.

5. A. B. Medvinsky, S. V. Petrovskii, I. A. Tikhonova, H. Malchow, B.-L. Li, SIAM review 44, 311 (2002).

6. J. van de Koppel, M. Rietkerk, N. Dankers, P. Herman, The American Naturalist 165, E66 (2005).

7. J. A. Commito, et al., Ecosphere 5, 1 (2014).

8. D. C. Thornton, European Journal of Phycology 37, 149 (2002).

9. E. J. Weerman, et al., The American Naturalist 176, E15 (2010).

10. S. Temmerman, et al., Journal of Geophysical Research: Earth Surface 110 (2005).

11. S. Fagherazzi, et al., Reviews of Geophysics 50 (2012).

12. T. van Der Heide, et al., Ecology 91, 362 (2010).

13. S. Mistr, D. Bercovici, Ecosystems 6, 61 (2003).

14. M. Bruschetti, Diversity 11, 168 (2019).

15. C. Boström, S. J. Pittman, C. Simenstad, R. T. Kneib, Marine Ecology Progress Series 427, 191 (2011).

16. D. B. Eggleston, L. L. Etherington, W. E. Elis, Journal of Experimental Marine Biology and Ecology 223, 111 (1998).

17. D. B. Eggleston, W. E. Elis, L. L. Etherington, C. P. Dahlgren, M. H. Posey, Journal of Experimental Marine Biology and Ecology 236, 107 (1999).

18. S. S. Bell, R. A. Brooks, B. D. Robbins, M. S. Fonseca, M. O. Hall, Biological Conservation 100, 115 (2001).

19. K. A. Hovel, H. M. Regan, Landscape Ecology 23, 75 (2008).

20. I. E. Gain, et al., Marine Biology 164, 8 (2017).

21. M. H. Hanke, M. H. Posey, T. D. Alphin, Marine Ecology Progress Series 581, 57 (2017).

22. L. Paquette, P. Archambault, F. Guichard, Marine Ecology Progress Series 608, 149 (2019).

23. L. Cornacchia, et al., Ecology 99, 832 (2018).

24. T. C. Chamberlin, Science 15, 92 (1890).

25. K. P. Burnham, D. R. Anderson, Springer-Verlag, New York, New York (2002).

26. D. R. Anderson, Model based inference in the life sciences: a primer on evidence (Springer Science & Business Media, 2008).

27. R Core Team, R: A Language and Environment for Statistical Computing, R Foundation for Statistical Computing, Vienna, Austria (2020).

28. Q. H. Vuong, Econometrica: Journal of the Econometric Society pp. 307–333 (1989).

29. M. D. Bertness, Ecology 65, 1794 (1984).

30. M. D. Bertness, S. D. Hacker, The American Naturalist 144, 363 (1994).

31. Q.-X. Liu, et al., Nature Communications 5, 5234 (2014).

